# Epitranscriptomic Reader YTHDF2 Regulates SEK1(*MAP2K4*)-JNK-cJUN Inflammatory Signaling in Astrocytes during Neurotoxic Stress

**DOI:** 10.1101/2024.01.26.577106

**Authors:** Emir Malovic, Alyssa Ealy, Phillip J. Hsu, Souvarish Sarkar, Cameron Miller, Dharmin Rokad, Cody Goeser, Aleah Kristen Hartman, Allen Zhu, Bharathi Palanisamy, Gary Zenitsky, Huajun Jin, Vellareddy Anantharam, Arthi Kanthasamy, Chuan He, Anumantha G. Kanthasamy

## Abstract

As the most abundant glial cells in the CNS, astrocytes dynamically respond to neurotoxic stress, however, the key molecular regulators controlling the inflammatory status of these sentinels during neurotoxic stress have remained elusive. Herein, we demonstrate that the m6A epitranscriptomic mRNA modification tightly regulates the pro-inflammatory functions of astrocytes. Specifically, the astrocytic neurotoxic stresser, manganese (Mn), downregulated the m6A reader YTHDF2 in human and mouse astrocyte cultures and in the mouse brain. Functionally, YTHDF2 knockdown augmented, while its overexpression dampened, neurotoxic stress induced proinflammatory response, suggesting YTHDF2 serves as a key upstream regulator of inflammatory responses in astrocytes. Mechnistically, YTHDF2 RIP-sequencing identified *MAP2K4* (*MKK4;* SEK1) mRNA as a YTHDF2 target influencing inflammatory signaling. Our target validation revealed Mn-exposed astrocytes mediates proinflammatory response by activating the phosphorylation of SEK1, JNK, and cJUN signaling. Collectively, YTHDF2 serves a key upstream ‘molecular switch’ controlling SEK1(*MAP2K4*)-JNK-cJUN proinflammatory signaling in astrocytes.

## Introduction

As the most abundant non-neuronal cells of the CNS, astrocytes are vital for brain and neuronal homeostasis. Their supportive functions include antioxidant defense, glutamate and water-ion-pH homeostasis, blood-brain barrier maintenance, synapse formation and maturation, and neurotrophic factor and cytokine production^1–4^. Astrocytes respond dynamically to neurotoxic stressors in order to protect and support neuronal health. When exposed to harmful substances or environments, astrocytes can undergo changes in their morphology and function, becoming reactive astrocytes. While this reactive response aims to limit damage and maintain homeostasis, chronic or sustained activation of astrocytes can also lead to uncontrolled inflammatory responses and contribute to various neurodegenerative conditions. Nevertheless, the precise molecular factors that govern the pro-inflammatory state of these guardian cells under chronic neurotoxic stress conditions remain enigmatic.

A substantial body of evidence has established that the neurotoxic concentrations of metal Mn significantly targets and adversely affects astrocyte physiology^5–9^. Indeed, astrocytes exhibit a significantly higher affinity for Mn than do neurons given their higher divalent metal transporter content^10,11^. Mn then accumulates through sequestration in mitochondria via the calcium uniporter^5,11,12^, exerting neurotoxic stress. Homeostatic dysregulation by Mn can induce inflammation in astrocytes, and chronic cellular alterations can transform quiescent astrocytes into reactive astrocytes, leading to neuronal toxicity^5,13–16^. In response to Mn, astrocytes release various chemokines, cytokines, and other neurotoxic factors and sustain these responses potentially through the NFκB and MAPK cascades^17,18^. Furthermore, chronic Mn exposure can induce Parkisonian conditions by primarily inducing astrocytic dysfunction^19,20^. Therefore, we adopted Mn treated astrocytes as model to understand the role of epitranscriptomic changes underlying chronic astrocyte activation

*N*^6^-methyladenosine (m6A) is the most prevalent epitranscriptomic modification^21^ regulating mRNA translation and decay. The DRACH (D = A/G/U, R = A/G, H = A/C/U) consensus sequences on mRNAs are targeted by the m6A writer complex, composed of METTL3 and METTL14 (methyltransferase-like), and accessory regulatory proteins such as WTAP, during which a methyl group is added to the sixth nitrogen position of adenosine^22–24^. Additionally, m6A modifications are reversible and can be removed by m6A demethylases such as ALKBH5 and FTO^25,26^. Recent studies reveal that m6A-modified mRNAs have reduced stability, yielding shorter half-lives that result in decreased time spent in ribosomal translation pools^27^. The m6A reader, YTHDF2 (YT521-B homology domain family), is known to promote the initiation of RNA decay; however, YTHDF1 has been defined to promote m6A mRNA translation, while YTHDF3 may facilitate both processes^28^. Understanding why such mechanisms would be requisite for cellular physiology and pathology varies across many biological disciplines, but the m6A readers which execute the fate of m6A mRNAs remain as fundamental elements of the ensemble. Since m6A modifications can dicate the fate of mRNA and translational rate, understanding its role in inflammatory processes would provide insights into upstream regulation of cytokine and chemokine mRNA production. In this regard, YTHDF2’s has been highlighted in certain cellular contexts, including hypoxia, survival/apoptosis, proliferation, and inflammation. In this regard, YTHDF2 has been shown to directly target secretory mRNAs such as IL-11^29^, transcription factor RELA, and MAP kinases^30–32^, revealing that YTHDF2’s reader function may be highly significant in inflammation. Pro-inflammatory astrocytic responses present with the upregulation of numerous chemokines and cytokines, which are regulated by multiple kinases and transcription factors^5,33,34^. However, the upstream epitranscriptomic regulatory events governing the neuroinflammatory signaling have yet to be delineated, hindering our ability to understand the molecular mechanism by which cytokine and chemokine mRNA dynamics are controlled during abberant astrocyte activation. To address this, herein, we adopted an approach using Mn as an astrocyte-specific neurotoxic stressor to determine whether m6A epitranscriptomic regulators modulate the neuroinflammatory response during astrocyte activation. Interestingly, our results reveal that epitranscriptomic reader YTHDF2 act as a negative epitranscriptomic regulator of the SEK1(*MAP2K4*)-JNK-cJUN proinflammatory signaling cascade in astrocytes, suggesting that YTHDF2 may be an exploitable target for controlling astrocytic inflammation in neuroinflammatory conditions.

## Results

### The astrocytic neurotoxic stressor Mn downregulates YTHDF2 expression in cell culture

As the major function of YTHDF2 in accelerating the decay of m6A mRNAs was recently demonstrated^27^, we reasoned this epitranscriptomic reader may influence the inflammatory signaling which relays rapid onset and decay depending on the insults. Only few studies have determined the expression levels of YTHDF2 under different stress conditions. Limited observations in cancer cells revealed heat-shock stress upregulated YTHDF2^35^, whereas hypoxia induced by oxygen deprivation or by cobalt chloride treatment downregulated YTHDF2^36^. Cellular hypoxia has been shown to precede oxidative stress responses^37^, and we recently reported that the astrocytic neurotoxic stressor Mn induced mitochondrial dysfunction and oxidative stress in astrocytes to augment neuroinflammation^5^. Previously, we showed that 100 μM Mn can evoke pro-inflammatory gene expression and the release of chemokines/cytokines in both primary mouse astrocytes and the human U251 astrocyte cells^5^, thus, we used 100 μM for all *in vitro* Mn treatment experiments. Primary mouse astrocytes isolated from 1-2 day old post-natal pups were treated with 100 μM Mn and revealed a time-dependent decrease in YTHDF2 protein levels beginning after 3 h and persisting across the entire 24 h period (Fig. 1A). Based on these results and previous studies showing the presence and sustainability of astrocytic inflammation, we continued using the 24 h Mn treatment timepoint^5^. Immunocytochemistry was also performed with primary mouse astrocytes, revealing primarily cytoplasmic perinuclear localization of YTHDF2 with notable decreases after Mn treatment (Fig. 1B). To test the specificity of Mn-induced down regulation of YTHDF2, wetreated human U251 astrocytes with individual metal chlorides (PbCl_2_, CuCl_2_, FeCl_2_) at 100 μM as well for 24 h and observed specific downregulation of YTHDF2 by Mn (Fig. 1C) but not with other metals. *YTHDF2* mRNA in U251 astrocytes showed small but significant decreases post Mn exposure, while *YTHDF1* and *YTHDF3* were not significant affected by Mn (Fig. 1D), suggesting Mn-induced YTHDF2 downregulation may be primarily attributed to the degradation of YTHDF2 protein. To understand how Mn may promote YTHDF2 degradation, we performed cotreatments of Mn and MG-132 (proteasome inhibitor) in U251 astrocytes and revealed inhibition of the proteasome prevents drastic decreases in YTHDF2 by Mn, as coindicated by increased amounts of the high molecular weight ubiquitinated proteins (Supplementary Fig. 1A), suggesting that Mn may promote YTHDF2 degradation via a proteasomal pathway such as the recently investigated SKP2-associated E3 ubiquitin ligase complex pathway^38^. Next, we investigated global m6A expression using LCMS/MS in U251 astrocytes. Since Mn decreases YTHDF2 protein levels, we hypothesized global m6A expression would increase; however, we observed a general decrease (Fig. 1E). These decreases in global m6A expression may be the result of increased m6A eraser FTO in U251s post Mn exposure (Supplementary Fig. 1B). Collectively, these results demonstrate Mn-induced neurotoxic stress downregulate a key epitranscriptomic reader YTHDF2 in astrocytic cells.

**Figure 1:**
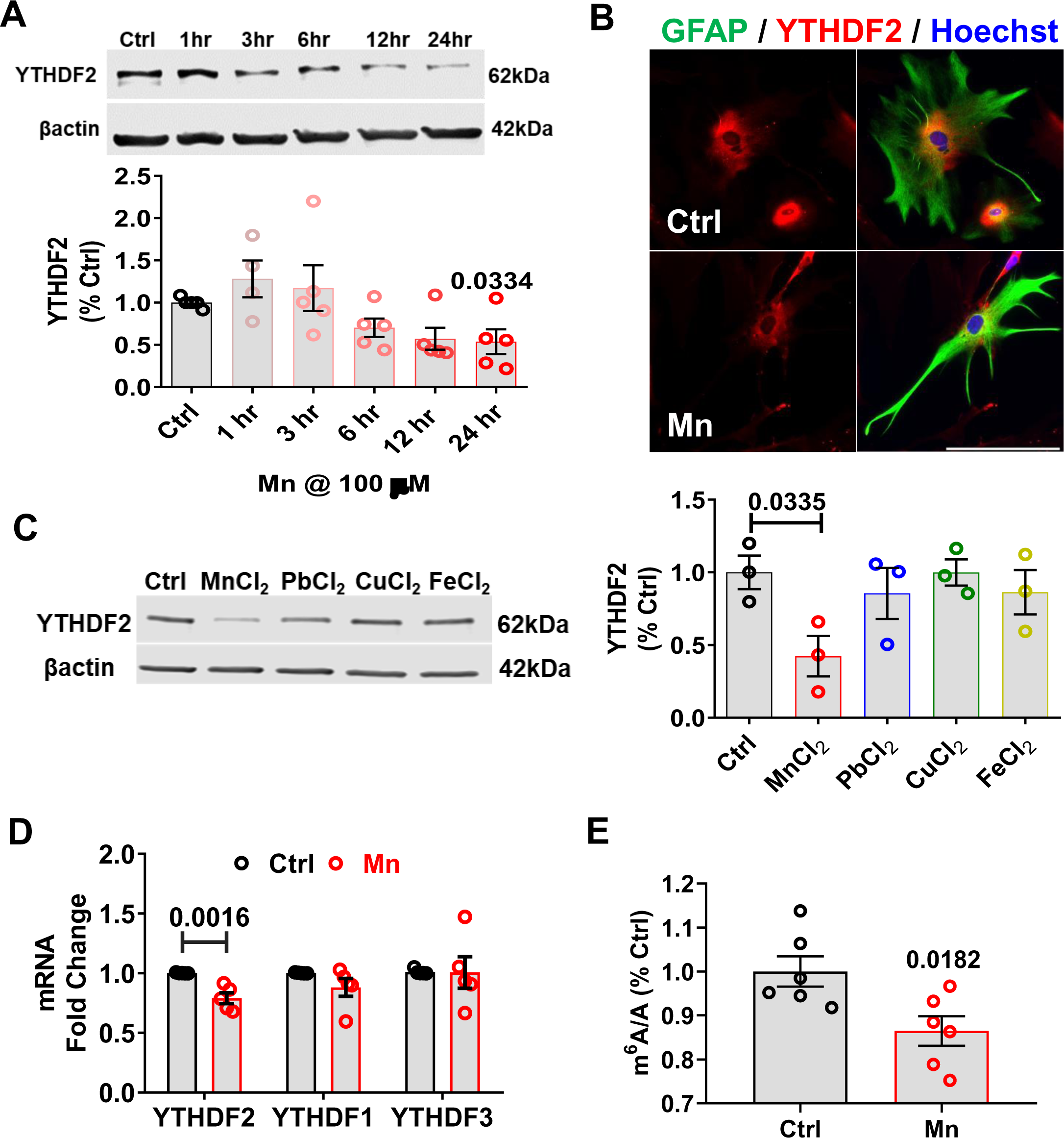
Mn treatment decreases YTHDF2 in cell culture. A) Primary mouse astrocytes were exposed to 100 μM for 1-24 h and YTHDF2 levels were determined by Western blot. The bottom panel show the results of densitometric analysis of YTHDF2 bands normalized by β-actin (n=4-5). YTHDF2 decreases time-dependently beginning after 3 h in primary mouse astrocytes. B) ICC representation at 40x depicting YTHDF2 decreases in Mn-exposed primary mouse astrocytes at 24 h. C) 24-hour treatment of human U251 astrocytes with different heavy metals, all at 100 μM (n=3). D) Mn decreases YTHDF2 mRNA at 24 hours in U251 astrocytes (n=5). E) Mn decreases global m6A levels at 24 hours in U251 astrocytes, as measured by LC-MS/MS (n=6). Data are means ± SEM. Two group comparisons performed using unpaired t-test. P-values ≤0.05 considered significant evidence.

### YTHDF2 levels can affect pro-inflammatory chemokine/cytokine responses in neurotoxic stress-exposed astrocytes

In response to neurotoxic stressors such as Mn, astrocytes activates pro-inflammatory signaling pathways that elicit the production of pro-inflammatory chemokines/cytokines^39^. The production of these pro-inflammatory mRNA transcripts could be increased due to prolonged half-lives in the ribosomal translating pools. Using SRAMP m6A site predictor^40^, we analyzed various proinflammatory transcripts for m6A DRACH sequences and found that most of them contain DRACH sequences within their coding sequences, suggesting m6A marks may be added by METTL3/METTL14 complex and read by the m6A reader proteins like YTHDF2. YTHDF2 can decrease mRNA stability through recruitment of CCR4-NOT^41^ or HRSP12–RNase P/MRP^42^ complexes, thus, we hypothesized the reduction of YTHDF2 may lead to an upregulated pro-inflammatory state by increased stability of pro-inflammatory transcripts, or indirectly by the stabilization of their upstream signaling factors. To test this hypothesis, we knocked down YTHDF2 transiently in U251 astrocytes using the *YTHDF2*-specific siRNA (si*YTHDF2*) for 48 h, followed by 24 h of 100 µM Mn treatment. Additionally, we also generated stably overexpressing YTHDF2-GFP U251 astrocytes to jointly determine whether Mn-induced YTHDF2 downregulation contributes to Mn-stimulated pro-inflammatory chemokine/cytokine responses. The specific knockdown and overexpression of YTHDF2, but not YTHDF2’s paralogs were validated by qRT-PCR and Western blot analyses (Fig. 2A-B). qPCR analysis revealed that siYTHDF2-transfected U251s had an overall pro-inflammatory basal state, which was exacerbated by Mn exposure, as indicated by *IL-1α*, *IL-1β*, *IL-8*, *IL-12α*, and *TNFα* (Fig. 2C). Conversely, overexpression of YTHDF2 showed no basal changes or reduced proinflammatory basal states, depending upon the chemokine/cytokine. Stable overexpression of YTHDF2 in U251 astrocytes markedly attenuated Mn-stimulated gene upregulation of the investigated chemokine/cytokine, except *IL-6* (Fig.2D). Interestingly, IL-6 showed similar patterns of gene expression when comparing YTHDF2 knockdown and overexpression states (Fig. 2C-D), suggesting YTHDF2 may have some degree of specificity, either directly upon certain chemokine/cytokines or indirectly upon certain upstream signaling factors, or maybe sensitive to exogenously driven genetic changes.

**Figure 2:**
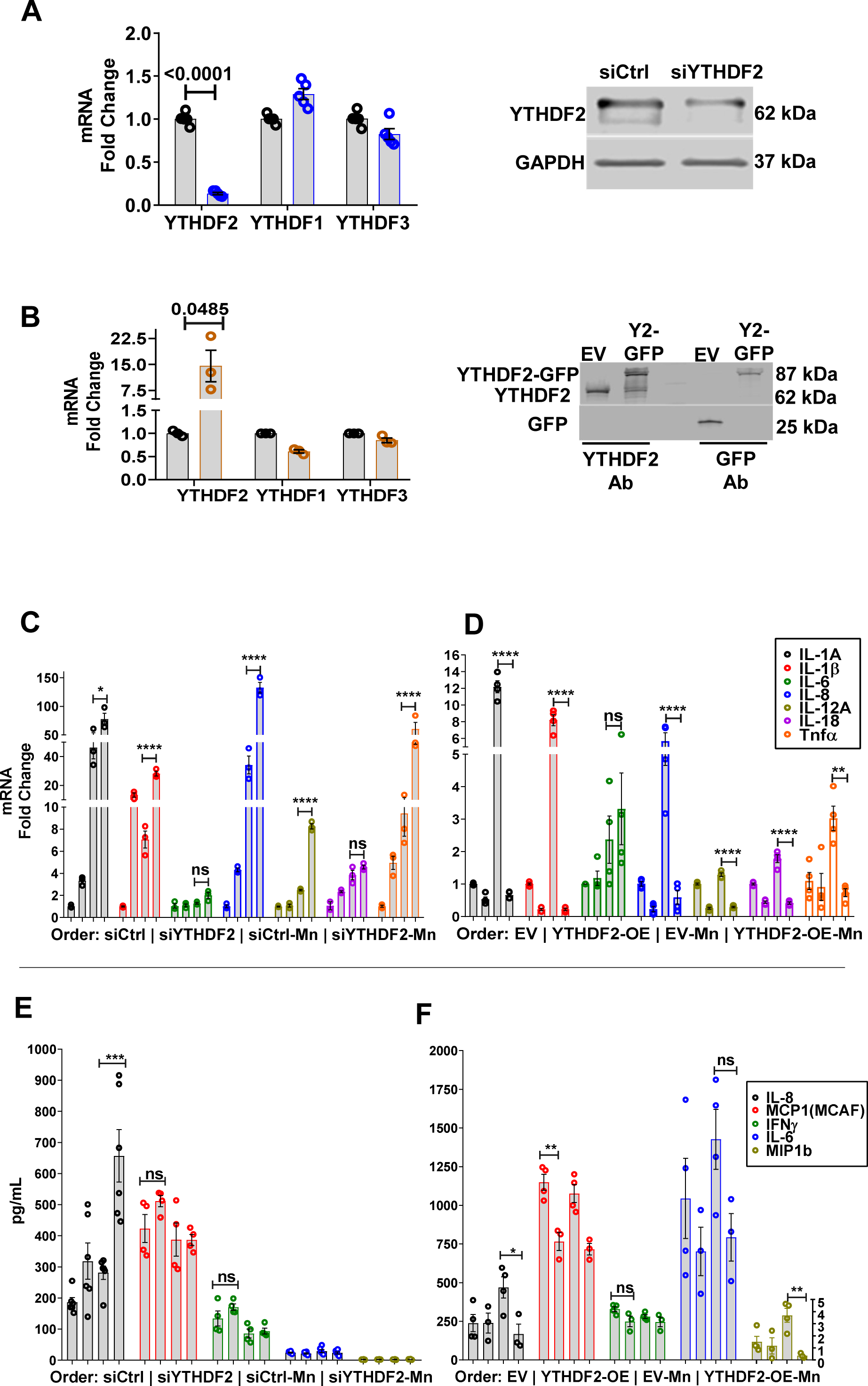
YTHDF2 levels can affect pro-inflammatory chemokine/cytokine responses in Mn exposed astrocytes. A) Validation of YTHDF2 knockdown using siRNA, both qPCR and immunoblotting (n=5-6). B) Validation of YTHDF2 overexpression, both qPCR and immunoblotting (n=3). C) siYTHDF2 increased basal pro-inflammatory gene expression and exacerbated after Mn exposure (n=3). D) Overexpression of YTHDF2 suppressed basal pro-inflammatory gene expression and prevented upregulation after Mn exposure (n=4). E-F) Mulitplex ELISA analysis of the treatment media of siYTHDF2 and YTHDF2 overexpression experiments for cytokines/chemokines (n=4-6). IL-8 was most consistently affected by YTHDF2 levels, showing exacerbated release in siYTHDF2 experiments, while its release was prevented in YTHDF2 overexpression experiments. MCP1 (MCAF), IFNγ, IL-6, and MIP1b showed similar trends but with less consistency overall. Data are means ± SEM. Two group comparisons performed using unpaired t-test. P-values ≤0.05 considered significant evidence. Two-way ANOVA with FDR Two-stage step-up method of Benjamini, Krieger and Yekutieli for multi-group comparison. Q-values ≤0.05 considered significant evidence.

Observations of differential gene expression of pro-inflammatory chemokine/cytokines upon knockdown and overexpression of YTHDF2 prompted us to assess if these same chemokine/cytokines were upregulated and released at the protein level. The treatment media from the gene expression experiments discussed above were assayed by the Luminex Bioplex ELISA system. Of the analyzed chemokine/cytokines, Mn exposure of U251 astrocytes had the greatest consistency on the secretion of IL-8. The knockdown of YTHDF2 elevated the basal level of IL-8 secretion. Upon Mn exposure, this response was significantly exacerbated as compared to siCtrl (siRNA Control) group exposed to Mn (Fig. 2E), corroborating the gene expression results for *IL-8* (Fig. 2C). MCP1, IFNγ, IL-6, and MIP1b remained largely unaffected or marginally variable after YTHDF2 knockdown and/or Mn treatment (Fig. 2E). Conversely, Mn-exposed EV (Empty Vector) astrocytes upregulated and secreted high levels of IL-8, while Mn-exposed YTHDF2-overexpressing cells did not upregulate and secrete IL-8 protein levels (Fig. 2F), corroborating the gene expression results (Fig. 2D). IL-6 and MIP1b responded similarly to IL-8, increasing in Mn-exposed EV cells but not in Mn-exposed YTHDF2-OE cells (Fig. 2F). MCP1 was significantly lower basally in YTHDF2-overexpressing cells, but Mn treatment did not affect MCP1 release in both stable cells (Fig. 2F). Corroborating the knockdown results (Fig. 2E), IFNγ release was not affected by YTHDF2 overexpression and/or Mn treatment (Fig. 2F). Additionally, we compared Mn-induced chemokine/cytokine levels to TNFα (100 ng/mL)-induced chemokine/cytokine levels, as TNFα is a potent pro-inflammatory cytokine (Supplementary Fig. 2A). Overall, we observed largely similar trends, especially when assessing YTHDF2 overexpressing cells. Notably, YTHDF2 overexpression significantly attenuated TNFα-induced levels of *IL-1β*, *IL-8*, *IFNγ*, and *MCP1*, while only *MCP1* approached statistical significance in YTHDF2 knockdown experiments (Supplementary Fig. 2A). These differences between Mn and TNFα in chemokine/cytokine induction may be the result of different signaling pathways or a difference in the potency of the two exogenous stimuli.

Based on our corroborated gene and protein results, we proceeded to perform an mRNA stability experiment using actinomycin D to determine if YTHDF2 directly regulates the mRNA decay of certain pro-inflammatory chemokine/cytokine genes. For this, we selected *IL-8* (based on Fig. 2C-D) and determined its mRNA stability in both YTHDF2 knockdown and overexpression U251 astrocytes (Supplementary Fig. 2B-C). Since basal levels of *IL-8* and other chemokine/cytokines were upregulated in siYTHDF2 cells, the stability of IL-8 mRNA in YTHDF2 knockdown cells was evaluated under non-stimulatory conditions. si*YTHDF2* cells showed no increased half-life of IL-8 over 2 and 4 h timepoints (Supplementary Fig. 2B). On the other hand, EV and YTHDF2 overexpression cells were evaluated under Mn exposure when the relative abundance of chemokines/cytokines is increased for an ease of comparison yet showed no sign of decreased half-life for *IL-8* mRNA (Supplementary Fig. 2C). These functional experiments suggest YTHDF2 have an anti-inflammatory function in astrocytes, as YTHDF2 overexpression attenuates astrocytic inflammation elicited by Mn. Since YTHDF2 does not appear to regulate IL-8 directly, it may be executing its effects upon the upstream regulators of pro-inflammatory pathways.

### YTHDF2 negatively regulates SEK1(MAP2K4)-JNK-cJUN pathway

YTHDF2 has been shown to target many mRNAs of signaling proteins involved in stress responses of varying conditions and cell types^30–32,43^. Since the actinomycin D mRNA stability assay showed no direct influence of YTHDF2 upon *IL-8* mRNA, we postulated that the pro-inflammatory response observed under the Mn exposure model may lie in the abundance of certain mRNAs that translate into signaling proteins or transcription factors whose downstream effects are to upregulate pro-inflammatory gene transcription. For that reason, we performed RNA-sequencing of Mn-exposed U251 astrocytes to evaluate the overall transcriptome in pursuit of corroborating the functional experiments of Figure 2. GO analysis of the Regulated Transcription Factor Targets revealed an overrepresentation of transcription factor families (bolded) including the AP-1 components (JUN and FOS) and HIF components (HIF1α and EPAS1) (Fig. 3A). To support, a network of significant differentially expressed genes (DEGs) was generated to provide a visual representation of direct and indirect gene regulation through transcription factors or protein-protein interactions (Supplementary Fig. 3A). In this DEG network, IL-1α and IL-8 were significantly upregulated genes by Mn exposure, supporting our functional results from Fig. 2, and further indicating their upstream regulation by the JNK pathway. Functional annotations of the main connected network of genes revolve around cell proliferation, apoptosis, cell adhesion, wound healing, angiogenesis, and responses to hypoxia. Overall, these RNA-sequencing results suggest Mn exposure can highly stress astrocytes leading to altered survival/apoptotic signaling, hypoxic signaling, and overall inflammatory reactivity.

**Figure 3:**
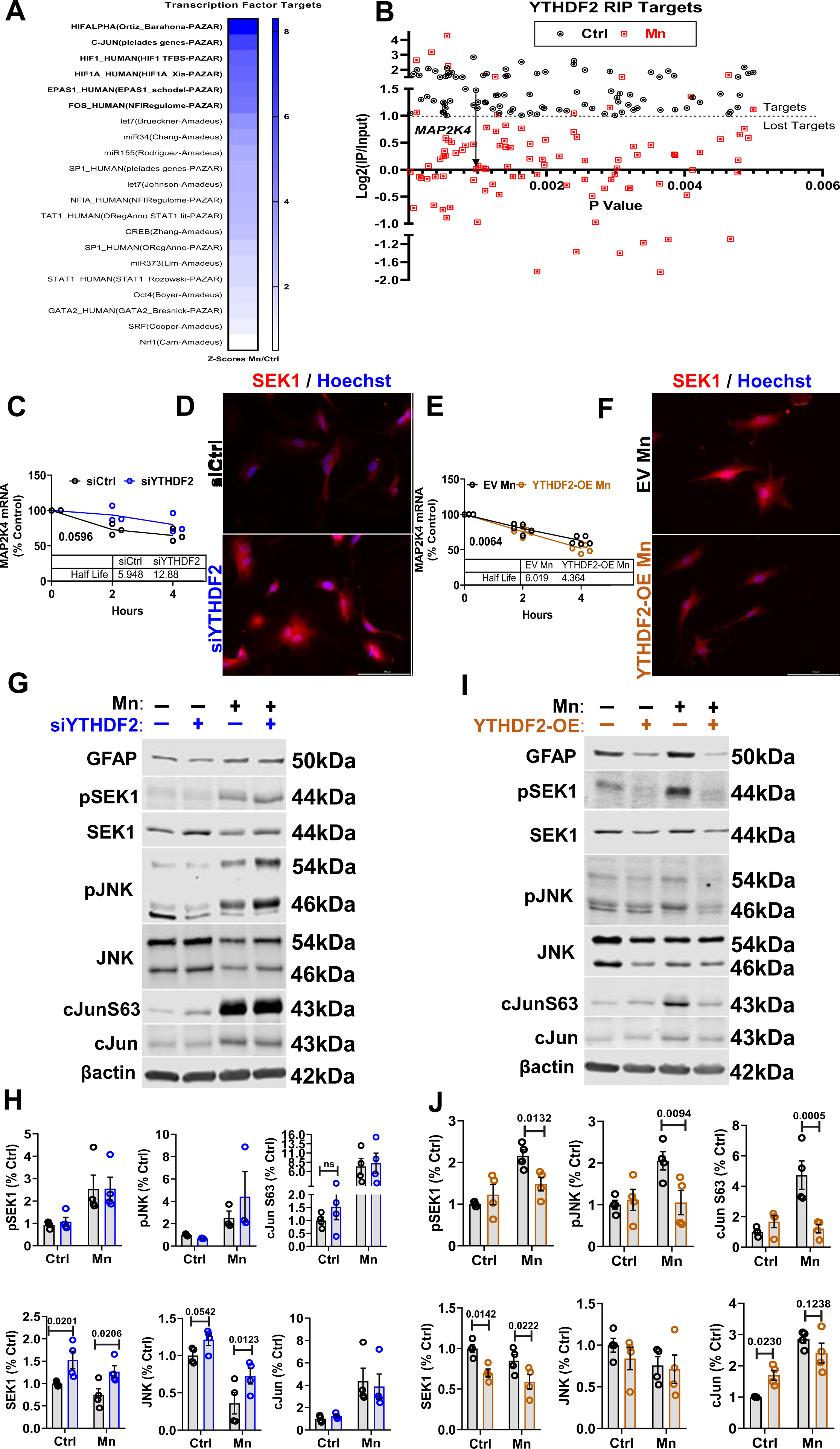
RNA- and RIP-sequencing of Mn exposed U251 astrocytes reveals YTHDF2 targeting of *MAP2K4* (SEK1). A) Z-scores of Mn/Ctrl GO for Transcriptional Factor Targets, indicating important roles for cJUN and HIF1α B) RIP-sequencing graphical representation of statistically significant YTHDF2 targets in Ctrl that were affected by Mn exposure. YTHDF2 targets are identified as having ≥+1 log2(RIP/input) ratio. *MAP2K4* was identified as a lost YTHDF2 target under Mn exposure, suggesting regulation of the SEK1(*MAP2K4*)-JNK-cJUN pathway. C) si*YTHDF2* astrocytes present with longer *MAP2K4* mRNA half-life (n=3). D) ICC representation at 40x depicting si*YTHDF2* astrocytes have increased SEK1 protein levels. E) YTHDF2 overexpressing astrocytes present with shorter *MAP2K4* half-life under Mn exposure (n=4-5). F) ICC representation at 40x depicting YTHDF2 overexpressing astrocytes have decreased SEK1 protein levels. G-H) Immunoblotting and quantification revealing cJUN phosphorylation is increased in Mn exposed astrocytes and sustained in siYTHDF2 Mn exposed astrocytes. SEK1 protein levels are basally increased in siYTHDF2 astrocytes (n=3-4). I-J) Immunoblotting and quantification revealing cJUN phosphorylation is increased in Mn exposed astrocytes and prevented in YTHDF2 overexpressing Mn exposed astrocytes. SEK1 protein levels are basally decreased in YTHDF2 overexpressing astrocytes (n=4). Data are means ± SEM. Two group comparisons performed using unpaired t-test, with 2-fold gene threshold and 1.96 Z-score threshold using Altanalyze. Adjusted P-values ≤0.05 considered significant evidence. For mRNA half-life comparisons, One Phase Decay Non-Linear Regression analysis was performed. P-values ≤0.05 considered significant evidence. Two-way ANOVA with FDR Two-stage step-up method of Benjamini, Krieger and Yekutieli for multi-group comparison. Q-values ≤0.05 considered significant evidence.

Given the strong hypoxic signature induced by Mn exposure in the RNA-sequencing data, we functionally validated it using si*HIF1α* in combination with Mn exposure. After siRNA transfection and Mn treatment, nuclear and cytosolic lysates were prepared to analyze HIF1α levels. Mn induced a strong upregulation of nuclear HIF1α, while si*HIF1α* transfection completely abolished HIF1α levels. Cytosolic HIF1α levels were undetectable likely because of rapid degradation and slow lysate processing methodology (Supplementary Fig. 4A). Interestingly, previous studies showed hypoxic states whether induced by O_2_ depletion^29^ or by cobalt chloride^36^ decrease YTHDF2 levels. In Zhong et al., YTHDF2 overexpression under hypoxia had a small effect in preventing HIF1α upregulation, suggesting YTHDF2 may target HIF1α mRNA^36^, providing one possible anti-inflammatory mechanism of controlling chemo/cytokine release, as HIF1α knockdown attenuated Mn-induced gene expression IL-1α and IL-1β (Supplementary Fig. 4B). Nuclear and cytosolic lysates of both siYTHDF2 and YTHDF2-OE cells revealed no observable increases in HIF1α in siYTHDF2-transefected cells, but there was an observable reduction in HIF1α levels in YTHDF2-OE cells (Supplementary Fig. 4C). Actinomycin D mRNA stability experiments did not support HIF1α mRNA as targets of YTHDF2 (data not shown). These results suggested YTHDF2 overexpression indirectly affects HIF1α protein levels, and HIF1α may only be partially responsible for the induction of certain chemokines/cytokines.

Next, we performed YTHDF2-RIP-sequencing assays on both control and Mn exposed U251 astrocytes to seek out plausible mRNA targets of YTHDF2 that, when degraded, would alleviate the pro-inflammatory response. Using the log2(RIP/input) calculation on our normalized TPM sequencing data, we denoted YTHDF2 targets as ≥+1.In brief, we identified a total of 1992 YTHDF2 transcript targets based on the control group. After performing a statistical comparison between control and Mn groups, we identified only 92 statistically significant (pvalue≤0.005) YTHDF2 targets, of which 90% were lost targets in the Mn group. Lost targets are defined as positive YTHDF2 targets in control conditions but are no longer significant YTHDF2 targets in the Mn group, putatively because of a decrease in YTHDF2 levels. Firstly, in agreement with our *IL-8* mRNA stability experiments, IL-8 was not an observed target of YTHDF2, nor were any other studied chemokines/cytokines. Secondly, HIF1α, ARNT, and EPAS1 were not observed as YTHDF2 targets either, eliminating the HIF components as candidates of YTHDF2 regulation of inflammation. And thirdly, as NFκB transcriptional activation has been shown to be important in regulating Mn toxicity and inflammation in astrocytes^17,44^ and as others have shown YTHDF2 to potentially regulate NFκB signaling^45^, we did seek out if NFκB or any other related components like RELA would be targets of YTHDF2. However, NFκB or its related components were not targets of YTHDF2 in our analysis. Nevertheless, our RIP-sequencing results did show that the mRNA of an upstream component of the MAPK pro-inflammatory signaling cascade, dual specificity mitogen-activated protein kinase kinase 4 (SEK1 (protein); *MAP2K4* or *MKK4* (gene)), was positively bound by YTHDF2 in the controls but presented as a lost target in Mn, suggesting *MAP2K4* mRNA is bound and directed towards decay by YTHDF2 (Fig. 3B). In response to various environmental stressors such as Mn, SEK1 may be activated through phosphorylation of its serine and threonine residues at positions 257 and 261. Activated SEK1 leads to phosphorylation of JNK, which further phosphorylates its main cellular substrate, cJUN. Our preliminary analysis of cJUN S63 revealed increased phosphorylation post-Mn treatment, and a sustained phosphorylation in si*YTHDF2* astrocytes treated with Mn. Conversely, YTHDF2 overexpression suppressed cJUN S63 phosphorylation (Supplementary Fig. 4D). The activated cJUN interacts with JunB, JunD, c-Fos or ATF to constitute the AP-1 transcription factor, which regulates gene expression that includes pro-inflammatory genes such as IL-8 and IL-1α^46–49^. A supplemental STRING analysis of SEK1, JNKs, JUN and its associated AP-1 partners, along with IL-8, demonstrated *MAP2K4*-JNK-cJUN signaling can lead to the transcriptional upregulation of IL-8 and IL-1α (Supplementary Fig. 4E), supporting YTHDF2’s indirect control of chemokine/cytokine mRNA and protein, which represents one mechanism of anti-inflammatory signaling in astrocytes.

To validate the YTHDF2 RIP-sequencing target *MAP2K4*, we first analyzed its mRNA sequence using the m6A prediction server, SRAMP^40^, and found a total of 12 DRACH consensus sites, 4 of which were very high confidence for m6A deposition (Supplementary Fig. 4F). Then, we performed functional mRNA stability experiments using actinomycin D to assess *MAP2K4*’s half-life in both YTHDF2 knockdown and overexpression cells. YTHDF2 knockdown led to a doubling of *MAP2K4*’s half-life over the timespan of 4 h (Fig. 3C). Since *MAP2K4*’s half-life was doubled, we performed immunocytochemistry to gauge whether *MAP2K4* protein (SEK1) was affected by the overall change in mRNA, and indeed, the abundance of SEK1 was also elevated (Fig. 3D). Conversely, in Mn-exposed, YTHDF2-overexpressing cells, we observed a significant decrease in the half-life of *MAP2K4* (Fig. 3E), which was also accompanied by a decrease of its protein SEK1’s immunoreactivity (Fig. 3F). These results support that *MAP2K4* transcripts can be destabilized by YTHDF2, which can overall influence the protein abundance of SEK1.

Since YTHDF2’s regulation of *MAP2K4* transcripts was potent enough to affect the abundance of its protein SEK1, we wanted to quantitatively determine if this mechanism affected the downstream phosphorylation and activation of cJUN, the principal component of the AP-1 transcription factor, which was an overrepresented transcription factor target within our RNA-sequencing dataset (Fig. 3A). SEK1 is the principal kinase responsible for promoting cJUN activation by first phosphorylating the JNK kinases, which then subsequently phosphorylate cJUN to trigger AP-1 transcription factor complex formation and activation that promote pro-inflammatory gene expression^50–53^. As such, we first assessed the signaling cascade under YTHDF2 knockdown conditions along with Mn exposure in U251 astrocytes. Under YTHDF2 knockdown only, SEK1 protein was upregulated, as observed previously by actinomycin D assay (Fig. 3C) and immunocytochemistry (Fig. 3D); however, there was no detectable changes in phosphorylation of SEK1. Under Mn exposure, phosphorylation of SEK1 was significantly upregulated and sustained in Mn-exposed YTHDF2 knockdown cells. Likewise, downstream partners JNK and cJUN were highly phosphorylated under Mn exposure and sustained in Mn-exposed YTHDF2 knockdown cells (Fig. 3G-H). Interestingly, YTHDF2 knockdown only cells had a notable elevation of cJUN phosphorylation observed by immunoblotting and immunocytochemistry(Fig. 3G-H, Supplementary Fig. 4D), suggesting that there may be some leaky basal elevation of the SEK1(*MAP2K4*)-JNK-cJUN cascade. Furthermore, total JNK protein levels behaved similarly to SEK1 protein levels; however, none of the JNK mRNAs (*MAPK8*, *MAPK9*, and *MAPK10*) were observed by YTHDF2-RIP-sequencing. Because cJUN phosphorylation was highly upregulated by Mn exposure and has been linked to biological processes such as inflammation, cell survival, and apoptosis, these results support that *MAP2K4* mRNA is a target of YTHDF2, and under YTHDF2 downregulation cJUN phosphorylation is sustained and saturated upon Mn exposure in astrocytes, leading to sustained pro-inflammatory signaling.

Lastly, we performed similar signaling experiments using YTHDF2 overexpressing cells to determine if Mn-induced SEK1, JNK, and cJUN phosphorylation can be prevented by YTHDF2 overexpressoion. Under unstimulated conditions, SEK1 protein was downregulated by YTHDF2 overexpression (Fig. 3I-J), supporting the findings from the mRNA stability assay and immunocytochemistry functional experiments (Fig. 3E-F). Again, there were no notable changes in phosphorylation of SEK1. Under Mn exposure, phosphorylation of SEK1 was significantly upregulated, but it was prevented in by YTHDF2 overexpression. Likewise, JNK and cJUN were highly phosphorylated in EV cells under Mn exposure, but the phosphorylation was significantly reduced in Mn-exposed YTHDF2-overexpressing cells (Fig. 3I-J). Interestingly, YTHDF2 overexpressing cells displayed a similar basal elevation of cJUN phosphorylation as observed in the YTHDF2 knockdown cells by immunoblotting (Fig. 3I-J) and immunocytochemistry (Supplementary Fig. 4D), suggesting that cJUN phosphorylation is sensitive upon disturbances in the abundance of YTHDF2. Collectively, these functional signaling studies support that astrocytes exposed to the neurotoxic stressor Mn upregulate the SEK1(*MAP2K4*)-JNK-cJUN signaling cascade by decreasing YTHDF2 levels to sustain proliferative, survival, and inflammatory signaling.

### YTHDF2 is decreased in Mn gavaged mice

We lastly examined YTHDF2 levels in mice gavaged with 30 mg/kg of Mn to verify our *in vitro* results. We evaluated the brain regions relevant to Mn neurotoxicity including the globus pallidus, the principal region affected in humans^54–57^, and the substantia nigra pars compacta/reticulata (SN), an additional site of interest where neuroinflammatory and neurotoxic effects have been observed^56–61^. In the globus pallidus, increased GFAP reactivity was prevalent in Mn-exposed mice (Fig. 4A). YTHDF2 immunoreactivity in GFAP-positive cells was generally observed to be decreased (Fig. 4A), despite the overall YTHDF2 immunoreactivity in GFAP-positive cells being low. Surprisingly, the predominant localization of YTHDF2 was in larger cytoplasmic cells, presumably neurons (Fig. 4A)^62^. We also immunoblotted whole tissue lysates from the substantia nigra and observed a general significant decrease in YTHDF2 levels, approximately ≤20% (Fig. 4B).

**Figure 4:**
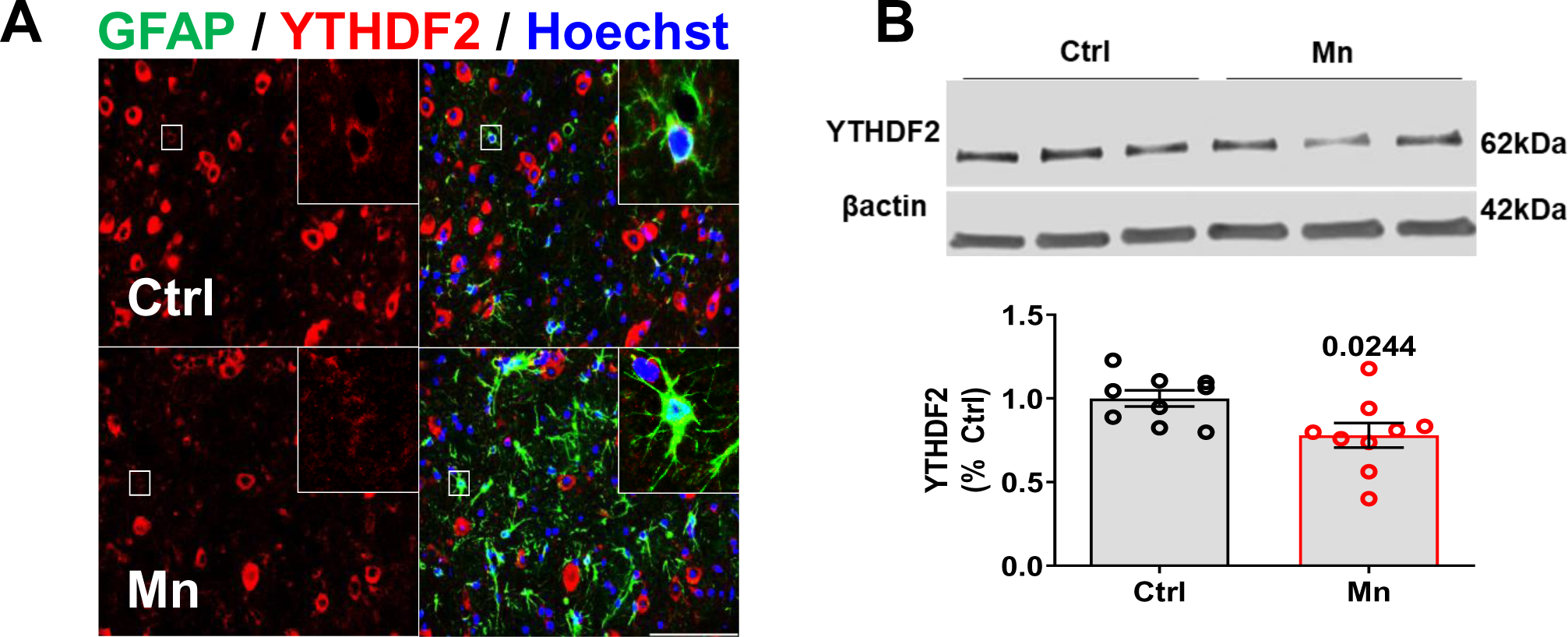
YTHDF2 is decreased in Mn gavaged mice. A) IHC representation at 40x depicting YTHDF2 decreases and colocalization in GFAP+ cells in the globus pallidus of Mn gavaged mice. B) Immunoblotting of the substantia nigra showing decreases of YTHDF2 in mice gavaged with Mn (n=9). Data are means ± SEM. Two group comparisons performed using unpaired t-test. P-values ≤0.05 considered significant evidence.

## Discussion

The overall destabilization effect m6A modifications confer upon RNAs, especially mRNAs that code for proteins, has promoted a wave of intense investigation in many disciplines such as cancer, virology, developmental biology, and immunology to name a few. Initial findings demonstrating specific m6A reader proteins are requisite for determining and executing the fate of m6A modified mRNAs provided this impetus^63^. YTHDF2 was shown to destabilize and promote the decay of m6A mRNAs^27^, whereas YTHDF1 could promote translation^64^, and YTHDF3 could potentially facilitate translation by association with YTHDF1 and afterwards associate with YTHDF2 to facilitate decay^28^. This putative mechanistic framework overall decreases the half-life of m6A mRNAs, specifically those modified within coding sequences or near 3’ UTR regions, suggesting m6A modifications may be necessary for critical cellular responses requiring large transcriptional bursts^21^. Under this framework, we postulated that YTHDF2 may be the most critical m6A reader as it principally promotes m6A mRNA decay. Herein, we show that YTHDF2 levels are decreased in astrocytes exposed to the astrocytic neurotoxic stressor Mn *in vitro* and in mice. Functional evaluation of YTHDF2 levels demonstrates that YTHDF2 negatively regulates *MAP2K4* (SEK1), thereby affecting the direct downstream JNK-cJUN signaling pathway, which can control survival, proliferation, and inflammation (Fig. 5). For the first time to our knowledge, decreased YTHDF2 levels allow astrocytes to mount and sustain a proper response to the environmental toxicant Mn.

**Figure 5:**
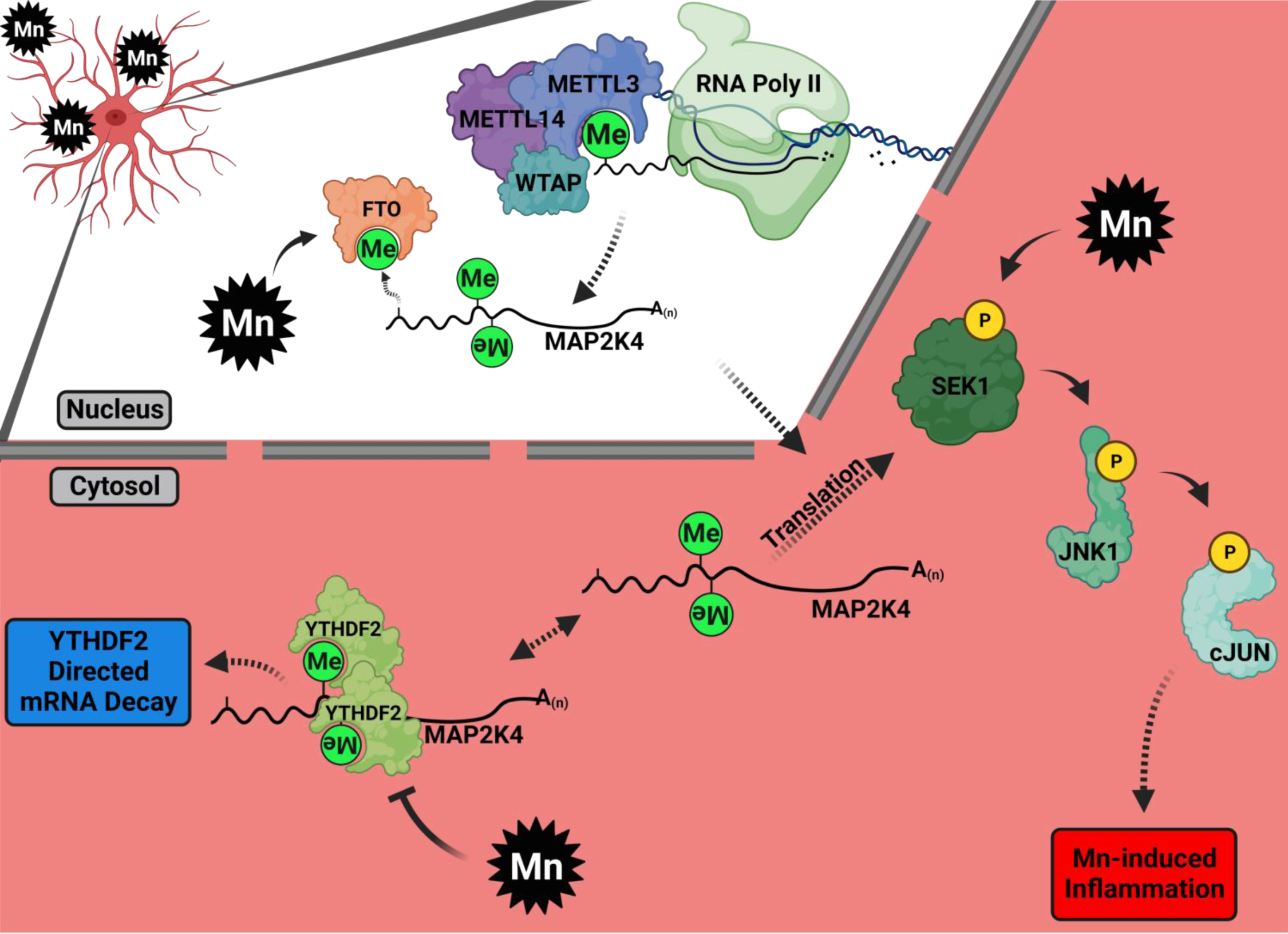
Integrated working hypothesis of YTHDF2’s role on the SEK1(*MAP2K4*)-JNK-cJUN pathway. Upon Mn exposure, FTO levels increase while YTHDF2 decreases, leading to increased half-life of *MAP2K4* mRNA. Increased abundance of *MAP2K4* leads to more SEK1 protein, allowing for the sustained activation (increased phosphorylation) of the downstream pathway targets JNK and cJUN by Mn, which promote the pro-inflammatory response in astrocytes. When YTHDF2 is overexpressed, *MAP2K4* mRNA is degraded. This leads to lower SEK1 protein levels, reducing the pathway activation of JNK and cJUN (decreased phosphorylation) by Mn, resulting an anti-inflammatory response.

The neurotoxic stressor Mn is a heavy metal known to perturb metabolic functions and redox reactions in all neural cells *in vitro*^5,58,65^. Additionally, it’s known to activate signaling pathways such as NFκB that can promote pro-inflammatory responses in astrocytes and microglia, which can inevitably damage neurons^17^. These observations support the importance of investigating neuroinflammatory responses as being major drivers to neurodegeneration in Mn neurotoxicity. Astrocytes are generally seen as maintainers of neural homeostasis, but in recent years more findings have surfaced demonstrating the importance of astrocytes in neuroinflammation, revealing their abilities to transform into neurotoxic cells^14,66^. These pro-inflammatory inductions require a large transcriptional burst of mRNAs necessary to elicit and sustain the response to combat the insults. As such, having mechanisms that could regulate mRNA turnover quickly becomes important in altering cellular states in response to environmental changes. In this study, we observed a time-dependent decrease in m6A reader YTHDF2 upon Mn exposure of astrocytes, in which levels declined after 3 h post Mn, being lowest at 12 and 24 h (Fig. 1A). Because Mn can elicit pro-inflammatory responses in astrocytes, this decrease in YTHDF2 levels would suggest that astrocytes need to decrease YTHDF2 to allow a sustained pro-inflammatory response. The mechanism by which Mn could decrease YTHDF2 levels is uncertain. Here we do observe small decreases in YTHDF2 transcripts, but the largest decrease appears to be post-translational, possibly through the proteasome as MG-132 prevented Mn-induced decreases of YTHDF2 (Supplementary Fig. 1A). Recently, it has been shown CDK1 can maintain the protein stability of YTHDF2, where inhibition of CDK1 led to high polyubquitination of YTHDF2 by the SKP2 E3 ubiquitin ligase complex^38^; however, it is unclear what effects Mn confers upon CDK1. Additionally, our results support the findings that hypoxia, whether induced by oxygen deprivation or cobalt chloride (CoCl_2_), can reduce YTHDF2 levels (Fig. 3A and Supplementary Fig. 4B) via an unidentified HIF1α mechanism^36^, or via HIF2α transcriptional repression^29^, both of which are increased in Mn-exposed astrocytes. With the decrease of YTHDF2 levels, we hypothesized an elevation in global m6A levels; however, the levels of m6A had decreased instead (Fig. 1E), signifying a more complex cellular response regarding m6A modifications and their associated regulatory tiers. High doses of arsenite have been shown to upregulate the m6A demethylases, while low doses did not^39^. Our dosage of Mn is considered a high dose^5,17^ and is in support of this finding. The demethylase, FTO, was upregulated by Mn treatment as shown by immunocytochemistry (Supplementary Fig. 1B). Such results present an interesting hypothesis concerning pro-inflammatory responses, whereby astrocytes and other cell types may elevate m6A demethylases and downregulate YTHDF2 concomitantly to ensure an increased half-life of mRNAs necessary to be translated for a given insult. These avenues remain to be fully elucidated, but what is certain is YTHDF2 overexpression can suppress pro-inflammatory gene expression and the production of specific chemokines/cytokines, of which IL-8 was most significantly affected in our experiments (Fig. 2C-F). This anti-inflammatory response of YTHDF2 is an agreement with other recent cancer studies showing suppression of IL-11 and IL-8^29,45^. Furthermore, although Mn failed to induce the production of MCP1 (also known as MCAF/CCL2) in our experiments, manipulation of YTHDF2 levels appreciably affected its basal levels, where the knockdown of YTHDF2 increased MCP1, and the overexpression of YTHD2 reduced MCP1 (Fig. 2E-F). Interestingly, TNFα significantly increased MCP1 levels that were exacerbated by YTHDF2 knockdown and suppressed by YTHDF2 overexpression (Supplementary Fig. 2A). This is a significant finding in that MCP1 has been shown to be a principal chemokine produced by astrocytes, even those stimulated by Mn^17,67^.

Our initial hypothesis was that YTHDF2 could bind various m6A mRNAs of secretory factors such as chemokines/cytokines, as in the case of IL-11 targeting in hepatocellular carcinoma^29^. YTHDF2-RIP-sequencing did not reveal any lost chemokine/cytokine targets by our methods. Others have previously shown that YTHDF2 can bind upstream signaling factors that lead to the production of pro-inflammatory chemokines/cytokines^30,32,45^. These signaling factors include RELA, *MAP2K4*, and MAP4K4. Our results pinpointed *MAP2K4* mRNA as a YTHDF2 target, lost under Mn exposure (Fig. 3B-J). *MAP2K4* or MKK4 (SEK1) is necessary for hepatocyte and fibroblast proliferation^51,68^ and for the protection against apoptosis in thymocytes^68^. Additionally, the SEK1(*MAP2K4*)-JNK-cJUN pathway has been shown to be upregulated in astrocytes upon infection with mutant retrovirus MoMuLV-ts1 leading to COX-2 upregulation^53^, and is upregulated in models of ischemia^69^. JNK phosphorylation is upregulated in *in vitro* astrocytes treated with lipopolysaccharide or cotreated with IFNγ^52,70^, and JNK expression is high in GFAP positive astrocytes in mechanical allodynia^71^. Indeed, in many stress models the SEK1(*MAP2K4*)-JNK-cJUN pathway is activated, as also observed in our Mn-exposed astrocytes (Fig. 3G-J; Supplementary Fig. 4D). YTHDF2 overexpression drastically prevented activation of the pathway by decreasing *MAP2K4* mRNA (Fig. 3C-F), thereby reducing total SEK1 levels and reducing phosphorylation of JNK and cJUN by Mn exposure. SEK1, JNK, and cJUN have all been found to be necessary for survival and proliferation^68,72,73^. Additionally, IL-8 and MCP1 are known targets of cJUN and the AP-1 transcription factor family, especially in astrocytes^74–76^, lending further explanatory power that would suggest these particular chemokines may be necessary for basal proliferation capabilities. Interestingly, GFAP levels were also reduced in YTHDF2 overexpressing cells (Fig. 3G & 3I), and GFAP is a known transcriptional target of cJUN^77^, providing another integrated explanation for the connection between the SEK1(*MAP2K4*)-JNK-cJUN pathway and motility^78^. Overall, YTHDF2 appears to significantly and negatively regulate the SEK-JNK-cJUN pathway that converges upon multiple cellular responses, which include survival/apoptosis, proliferation, migration, and inflammation.

Our observations of the negative correlation between YTHDF2 levels and Mn-induction of pro-inflammatory genes/proteins in astrocytes suggested YTHDF2 may act as anti-inflammatory m6A reader by negatively regulating the SEK1(*MAP2K4*)-JNK-cJUN pathway. In our examination of mice gavaged daily with Mn at 30 mg/kg for 30 days, YTHDF2 levels showed evidence of modest decreases in the substantia nigra and in astrocytes of the globus pallidus (Fig. 4A-B). However, we observed no significant neuronal death, suggesting higher doses of Mn and/or longer timepoints in Mn gavage studies are necessary to see if they are more effective in reducing YTHDF2. Alternatively, other Mn administration models should be tried to establish consistency and efficacy for disease modeling, especially inhalation models. Toxicity from inhaled Mn was first observed in Mn ore crushers^79,80^. It is observably more potent based on a recent study nasally administering 30 mg/kg for 21 days, in which dopaminergic cell (tyrosine hydroxylase +) loss was evident in the substantia nigra^81^; however, the globus pallidus was not investigated.

## Conclusion

In summary, our studies indicate that YTHDF2 targets *MAP2K4* mRNA to negatively regulate the SEK1(*MAP2K4*)-JNK-cJUN pathway in astrocytes, which is known to converge upon survival/apoptotic, proliferative, migratory, and inflammatory responses. Furthermore, m6A perturbation and decreases in YTHDF2 in our animal model of Mn neurotoxicity support the relevance of m6A epitranscriptomics in other neuroinflammatory and neurodegenerative conditions^82^. Overall, YTHDF2’s m6A reader roles may be exploited for therapeutic benefits in controlling processes underlying chronic neurodegenerative diseases.

## MATERIALS AND METHODS

### Chemicals & Reagents

Cell culture media, supplements, and transfection reagents (lipofectamine 2000 in conjunction with Opti-MEM) were purchased from Life Technologies (Waltham, MA). SAFC fetal bovine serum (FBS), manganese chloride tetrahydrate (MnCl_2_), and puromycin were obtained from Sigma (St. Louis, MO), while lead chloride (PbCl_2_), copper chloride (CuCl_2_), and iron chloride (FeCl_2_) were purchased from Fisher Scientific (Waltham, MA). The proteasome inhibitor MG-132 was purchased from Tocris Bioscience (Minneapolis, MN). Antibodies used are as follows: rabbit YTHDF2 (Proteintech, Rosemont, IL: #24744-1-AP), mouse GFAP (EMD Bioscience, Burlington, MA: #MAB360), rabbit GFAP (EMD Bioscience: #AB5804), rabbit FTO (Novus Biologicals: #NB110-60935), mouse HIF1α (BD Biosciences, Becton Drive Franklin Lakes, NJ: #610959), rabbit Lamin B1 (Abcam, Cambridge, UK: #ab16048), rabbit pSEK1 (Cell Signaling Technology (CST), Danvers, MA: #9156), rabbit SEK1 (CST: #9152), rabbit cJUN S63 (CST: #2361), mouse cJUN (Santa Cruz, Dallas, TX: #sc-74543), rabbit pJNK (CST: #4671), rabbit JNK (CST: #9258), mouse β-actin (Sigma: #A5441), rabbit β-actin (Sigma: #A2103), mouse GAPDH (Sigma: #G8795), and goat GFP (Abcam: ab6673).

### Cell culture, Treatments, & Transfections

For primary mouse astrocytes, 1-2-day-old mouse pups (C57BL/6) were decapitated, and their brains harvested. Subsequently, the meninges were removed, and the brains were incubated in 0.25% Trypsin-EDTA for 15 min in a 37°C water bath and mixed every 5 minutes. Afterwards the brains were washed in DMEM/F12 growth media supplemented with 10% FBS, 1% penicillin/streptomycin, 1% L-glutamine, 1% sodium pyruvate, and 1% non-essential amino acids, homogenized by pipetting, and filtered through a 70-μm filter. The resulting cell suspension was then plated in flasks to attach and grow for 16 d. The growth media was aspirated and replaced with fresh growth media on the 6^th^ d. After 16 d, the cells were collected and subjected to CD11b-positive selection to remove microglia. The negative fraction containing astrocytes was maintained in DMEM media supplemented with 10% FBS, 1% glutamine, and 1% penicillin/streptomycin, and treated with metals in DMEM containing 2% FBS.

The human U251-MG astrocyte cell line obtained from ATCC (HTB-17) was maintained in MEM supplemented with 10% FBS and 1% penicillin/streptomycin and treated in MEM with 2% FBS. To generate a stable cell line expressing the human YTHDF2, U251-MG cells were stably transfected with a GFP-tagged YTHDF2 expression plasmid pLenti-C-mGFP-P2A-Puro (OriGene, #RC230306L4) or GFP empty vector by lipofectamine 2000 according to the procedure recommended by the manufacturer. The stable transfectants were selected and maintained in 3 μg/mL of puromycin added to the growth media.

The siRNA transfection of U251 astrocytes was performed by using lipofectamine 2000 following the manufacturer’s instructions and scaled as necessary with a default of 100 pmol siRNA for a 10 cm^2^ area. The YTHDF2-specific siRNA and AllStars negative control siRNA (#1027281) were purchased from Qiagen, while the HIF1α-specific siRNA was obtained from Ambion. The siRNA sequence for YTHDF2 is 5′-AAGGACGTTCCCAATAGCCAA-3′ and for HIF1α is 5’-GCTGATTTGTGAACCCAT-3’ (Ambion). After 48 h from the initial transfection, the cells were subjected to Mn treatment experiments. For all cell experiments, MnCl_2_ treatments were performed at 100 μM, and likewise, so were PbCl_2_, CuCl_2_, and FeCl_2_ treatments. Treatment with the proteasome inhibitor MG-132 was performed at 5 μM. Actinomycin D treatments were performed at 10 μg/mL.

### Animal Studies

All animal procedures were approved by Iowa State University’s Institutional Animal Care and Use Committee (IACUC). Mice were housed under standard conditions for constant temperature (22 ± 1°C), relative humidity (30%), and a 12-h light cycle with food and water available ad libitum. Eight-week-old C57BL/6NCrl mice purchased from Charles River were orally gavaged with Mn (30 mg/kg) dissolved in ultrapure water daily for 30 days. This dose was chosen based on previous studies on mouse models of Mn neurotoxicity, which generally use the Mn doses between 5-30 mg/kg/day for periods of 21-120 days^83^. The control animals received daily injections of the same volume of water.

### Immunocytochemistry (ICC) and Immunohistochemistry (IHC)

For ICC, 13-mm coverslips were placed into 24-well plates and coated with poly-D-lysine. After experimental treatments, the media was aspirated, and 4% paraformaldehyde (PFA) was added to the wells, which were then incubated for 30 min at room temperature. Afterwards, the coverslips were rinsed 5 times with PBS and then blocked using 5% donkey serum with 0.25% triton X-100 in PBS for 1 h at room temperature. Primary antibodies were incubated overnight at room temperature, while secondary AlexaFluor antibodies, all from Invitrogen, were incubated for 1 h at room temperature. Washing was performed using PBS. Coverslips were then imaged using a Keyence All-in-one microscope (Keyence, Osaka, Japan).

For IHC, mice were perfused with 4% PFA, after which the brains were placed in 4% PFA for at least 24 h. For paraffin-embedded sectioning, the brains were placed onto a matrix and cut into 2-mm coronal pieces and then embedded into cassettes. Paraffin-embedded sections were cut at 5 μm. The sections were rinsed in PBS and then subjected to antigen retrieval solution (10 mM sodium citrate, pH 7.6). The sections were blocked using 5% donkey serum with 0.25% triton X-100 in PBS for 1 h at room temperature. Primary antibodies were incubated overnight, while secondary antibodies were incubated for 1 h at room temperature. The sections were imaged using the same Keyence All-in-one microscope.

### Immunoblotting

Cells or brain tissues were homogenized in modified RIPA buffer containing Halt protease and phosphatase inhibitors, 1 mM sodium orthovanadate, and 1 mM phenylmethanesulfonyl fluoride. Afterwards, the lysate was sonicated, cleared by centrifugation at 4° C, and the supernatant was collected. The protein concentrations were determined by a Bradford assay. Ten to forty μg of protein was loaded for SDS-PAGE gel electrophoresis and transferred on to nitrocellulose membranes. Membranes were blocked using Li-cor Intercept blocking buffer for 1 h at room temperature. Primary antibodies were incubated overnight at room temperature, while secondary antibodies were incubated for 1 h at room temperature. Washing was performed using PBS-Tween. All membranes were imaged using an Li-cor Odyssey infrared imaging system (Li-cor, Lincoln, NE), and data were analyzed using ImageJ software or Odyssey software 2.0 (Li-cor). β-actin or GAPDH served as an internal control for loading.

### Nuclear and Cytoplasmic Fractionation

Cells were collected by scraping and washing with PBS 1 time. Twenty μL of the packed cell volume was used with CER I (200 μL), CER II (11 μL), and NER (100 μL) buffers according to manufacturer’s protocol of the NE-PER™ Nuclear and Cytoplasmic Extraction Reagent kit (Thermo Fisher Scientific). Afterwards, the immunoblotting procedure given above was followed, with 10 μg of protein from both the nuclear and cytosolic fractions loaded onto the same gel.

### Luminex ELISA Assays

For chemokine/cytokine analysis, the treatment media was collected and analyzed using the Bio-Plex Pro Human Cytokine 17-plex Assay kit (#M5000031YV) following the established kit protocol. The analytes were detected using the Luminex Bio-Plex 200 system and quantified against a known standard curve, which is represented in picograms/milliliter (pg/mL).

### Quantitative real-time RT-PCR (qRT-PCR)

To extract RNA, cells were lysed using 1 mL of TRIzol reagent. Chloroform (200 μL) was added, and samples were vigorously mixed for 10 sec, allowed to settle for 3 min at room temperature, and then centrifuged for 15 min at 12,000 rcf 4° C to obtain phase separation. The top aqueous layer was collected, mixed with isopropanol (at least 1:1), incubated for 10 min at room temperature, and then centrifuged for 15 min at 12,000 rcf 4° C. Isopropanol was removed, while the RNA pellets were washed with 75% ethanol and then centrifuged for 15 min at 20,627 rcf 4° C. After air-drying for 10 min at room temperature, the RNA pellets were dissolved in RNase/DNase free ultrapure water and quantified using a NanoDrop 2000 spectrophotometer (Thermo Fisher Scientific). One microgram of RNA was converted into cDNA using a High-Capacity cDNA Reverse Transcription kit (Applied Biosystems). qPCR was performed using PowerUp Sybr Green reagents (Applied Biosystems) and run on the QuantStudio 3 system (Applied Biosystems, Waltham, Massachusetts). Human *18S* rRNA was used as the housekeeping gene for normalization. The primer sequences for the *YTHDF* paralogs and *MAP2K4* are as follows (5’-3’): *YTHDF1* (F: ACACAACCTCCATCTTCGAC & R: ACTGGTTCGCCCTCATTG), *YTHDF2* (F: TAGCCAACTGCGACACATTC & R: CACGACCTTGACGTTCCTTT), *YTHDF3* (F: TGACAACAAACCGGTTACCA & TGTTTCTATTTCTCTCCCTACGC), and *MAP2K4* (F: TCCCAATCCTACAGGAGTTCAA & R: CCAGTGTTGTTCAGGGGAGA). Validated QuantiTect primer sets obtained from (Qiagen) for *IL-1α* (QT00001127), *IL-1β* (QT00021385), *IL-8* or *CXCL8* (QT00000322), *IL-12α* (QT00000357), *IL-18* (QT00014560), *TNFα* (QT00029162) were also used. The data were analyzed using the ΔΔC_t_ method.

### Actinomycin D mRNA Stability Assay

After transfection and/or treatments, cells were treated with 10 μg/mL of actinomycin D (Invitrogen) for 2 and 4 h, with 0 h as the initial condition. Cells were then collected by scraping and washing with PBS once. RNA was isolated and quantified as described in the qRT-PCR section.

### RNA- and RIP-sequencing and Analysis

The RNA- and RIP-sequencing analysis was adapted from previous studies^27,41^. Briefly, cells were lysed in 2 volumes of modified TNE buffer containing 1% NP-40, Halt protease and phosphatase inhibitors, 1 mM sodium orthovanadate, 1 mM phenylmethanesulfonyl fluoride, and SUPERase RNase inhibitor (Invitrogen, 1:100). Lysates were cleared twice through centrifugation, and 2.5% of the total cleared lysate was collected and mixed with 1 mL of TRIzol for the RNA input fraction. RNA input was poly (A) purified using the GenElute mRNA kit (Sigma, #MRN 70). The remaining cleared lysate was subjected to immunoprecipitation with 10 μg of YTHDF2 antibody under constant rotation at 4° C for 4 h. The Protein A/G Magnetic Beads from Thermo Scientific Pierce (#88803) were then added to incubate under constant rotation at 4° C for 1 h. Then the YTHDF2 mRNP complexes were magnetically isolated, washed with TNE buffer, and placed in 1 mL of TRIzol for RNA isolation for the immunoprecipitated fraction. Sequencing was performed using a 50 cycle (single-end) with a HiSeq 3000 system (Illumina) with a total 100 ng of RNA from both input and immunoprecipitated fractions. Fastq files were processed with the default settings of Altanalyze (embedded with Kallisto), and the expression files were subsequently processed through the embedded gene ontology software GO-Elite^84–86^. For RIP-seq targets, we used the log2(RIP/input) calculation on our TPM sequencing data, with values ≥+1 as denoted as targets of YTHDF2, while values <+1 were considered non-targets of YTHDF2^87^.

### Global m6A LC-MS/MS

Global m6A was quantified as previously described^27^. Briefly, RNA was isolated by TRIzol and subjected to 2 rounds of poly (A) purification using the Gen-Elute mRNA kit (MRN 70). Thirty nanograms of mRNA were digested by nuclease P1 at 37 °C for 3 h, followed by FastAP alkaline phosphatase digestion at 37 °C for 6 h. The samples were diluted and filtered through a 0.22-μm filter, and 5 µl of each sample was injected into the LC-MS/MS. Reverse phase UPLC using a C18 column separated the nucleosides, which were detected by an Agilent 6410 QQQ LC mass spectrometer in positive electrospray ionization mode. Quantification was performed using a standard curve after taking the nucleoside to base ion mass transitions of 282 to 150 (m6A), and 268 to 136 (A). The ratio of m6A to A was calculated and expressed as a percentage of the control.

### Statistical Analysis

GraphPad 8.0 was utilized for data visualization and statistical analysis. For two group comparisons, we applied the unpaired t-test. Welch correction was applied for comparisons with unequal variance. For multiple group comparisons, two-way ANOVA was applied, with corrections for multiple comparisons controlled using FDR two-stage step-up method of Benjamini, Krieger and Yekutieli. For mRNA half-life comparisons, one-phase decay non-linear regression analysis was performed. P-values less than 0.05 were considered as strong evidence; however, p-values between 0.05 and 0.01 were considered as weak evidence taking into consideration the variations in data and trends^88^.

## ACKNOWLEDGMENTS

This work was supported by the National Institutes of Health (NIH) R01 Grants ES010586, ES027245, ES026892, ES034196, NS100090, NS121692, (AGK) and NS124226 (AK). In addition, AGK was also supported by the W. Eugene and Linda Lloyd Endowed Chair, the Scott and Nancy Armbrust Biomedical Science Endowment, the Johnny Isakson Endowed Chair, and the Georgia Research Alliance Eminent Scholar funds.

## AUTHOR CONTRIBUTIONS

EM and AE contributed equally in conceiving, performing experiments, and writing and revising the manuscript as necessary. PJH and SS provided insights and performed experiments. DR, CG, AKH, AZ, and BP performed experiments. GZ, HJ, VA, and AK provided critical feedback for the work and performed manuscript revisions. CH provided resources for performing RNA- and RIP-sequencing and the initial *Ythdf2*-floxed mice. AGK conceived, provided critical feedback, manuscript revisions, and general guidance.

