## Supplemental Figs.1-4 for "Epitranscriptomic Reader YTHDF2 Regulates SEK1(*MAP2K4*)-JNK-cJUN Inflammatory Signaling in Astrocytes during Neurotoxic Stress"

### SUPPLEMENTARY INFORMATION

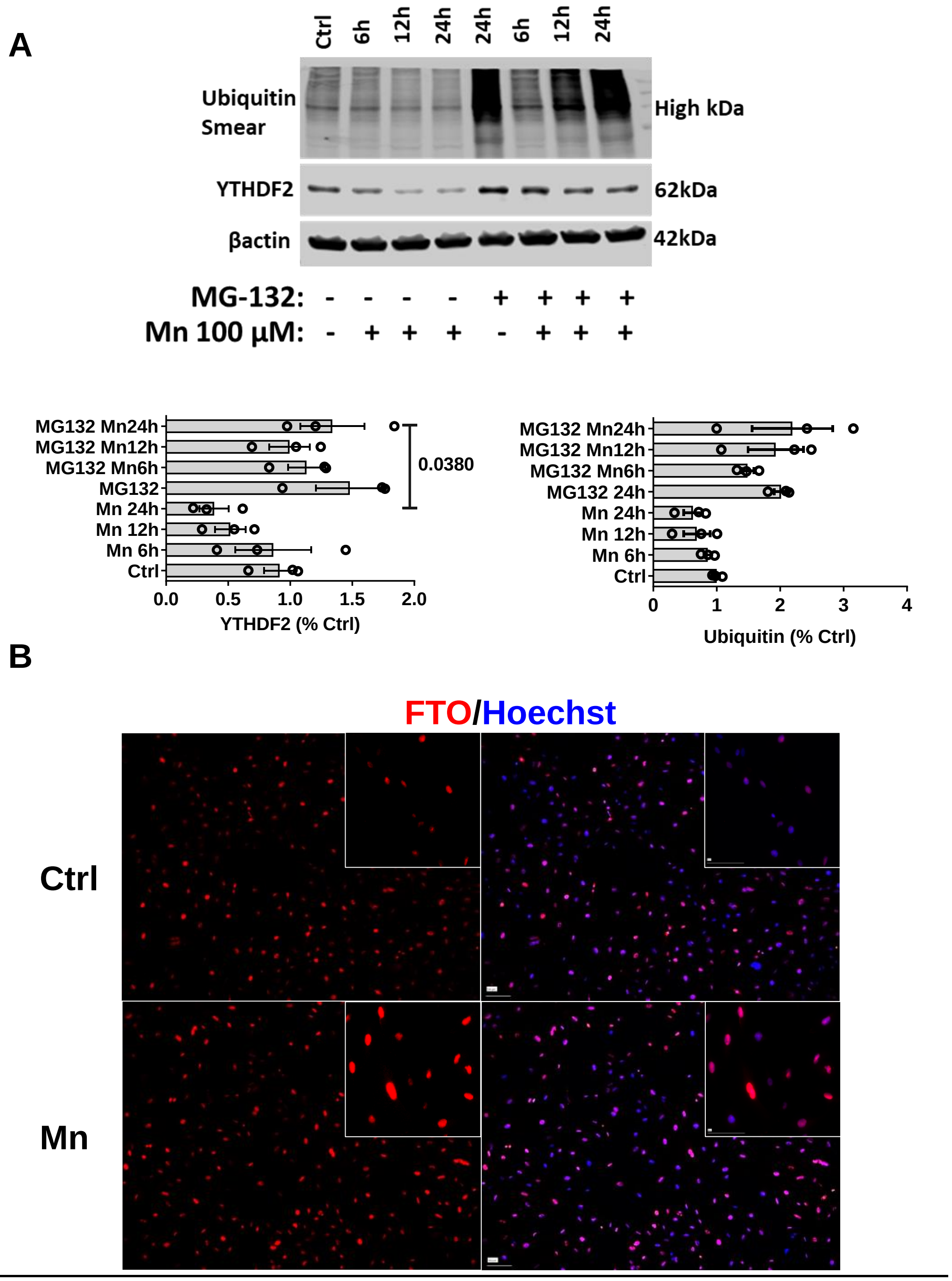


**Supplementary Figure 1: YTHDF2, ubiquitin, and FTO levels in Mn exposed astrocytes.** A) Proteasomal inhibitor, MG-132, prevents loss of YTHDF2 during co-exposure with Mn (n=3). B) ICC representation at 4x and 40x (inset) depicting increases in m6A eraser, FTO. Data are means ± SEM. One-way ANOVA with FDR Two-stage step-up method of Benjamini, Krieger and Yekutieli comparison. Q-values ≤0.05 considered significant evidence.

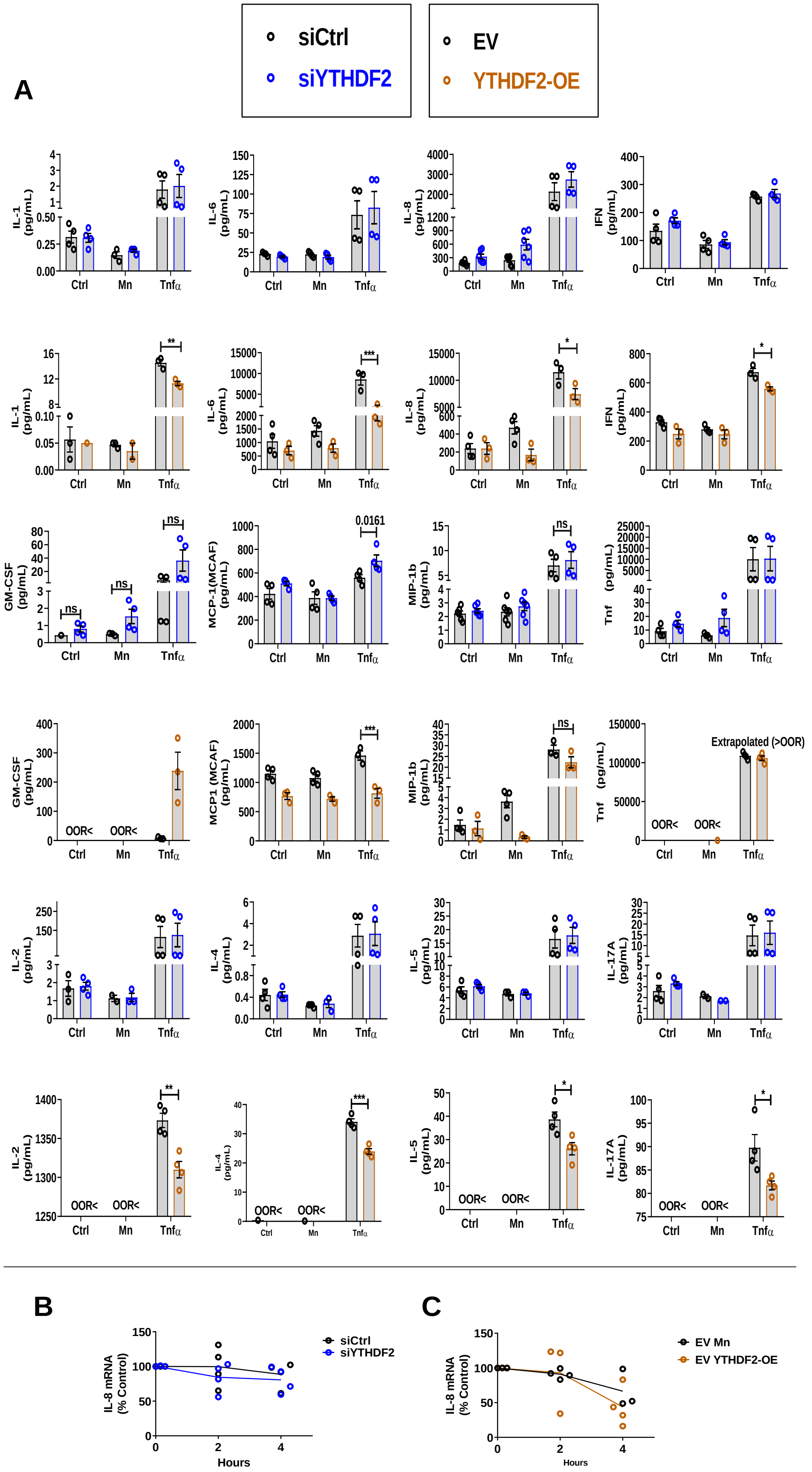
A
siCtrl
siYTHDF2
EV
YTHDF2-OE
IL-1 (pg/mL)
IL-6 (pg/mL)
IL-8 (pg/mL)
IFN (pg/mL)
GM-CSF (pg/mL)
MCP-1(MCAF) (pg/mL)
MIP-1b (pg/mL)
Tnf (pg/mL)
MCP1 (MCAF) (pg/mL)
IL-2 (pg/mL)
IL-4 (pg/mL)
IL-5 (pg/mL)
IL-17A (pg/mL)
Ctrl
Mn
Tnfα
ns
0.0161
Extrapolated (>OOR)
OOR<
B
IL-8 mRNA (% Control)
Hours
siCtrl
siYTHDF2
C
EV Mn
EV YTHDF2-OE

**Supplementary Figure 2: Multiplex ELISA assay for chemokines/cytokines in siYTHDF2 and YTHDF2 overexpression, along with TNFα comparison in astrocytes.** A) TNFα exposure showing similar effects as Mn exposure (n=3-4). B) Actinomycin D mRNA stability assay for IL-8, in which IL-8 does not appear to be a direct of YTHDF2 (n=3-4). Data are means ± SEM. Two-way ANOVA with FDR Two-stage step-up method of Benjamini, Krieger and Yekutieli for multi-group comparison . Q-values ≤0.05 considered significant evidence. For mRNA half-life comparisons, One Phase Decay Non-Linear Regression analysis was performed. P-values ≤0.05 considered significant evidence.

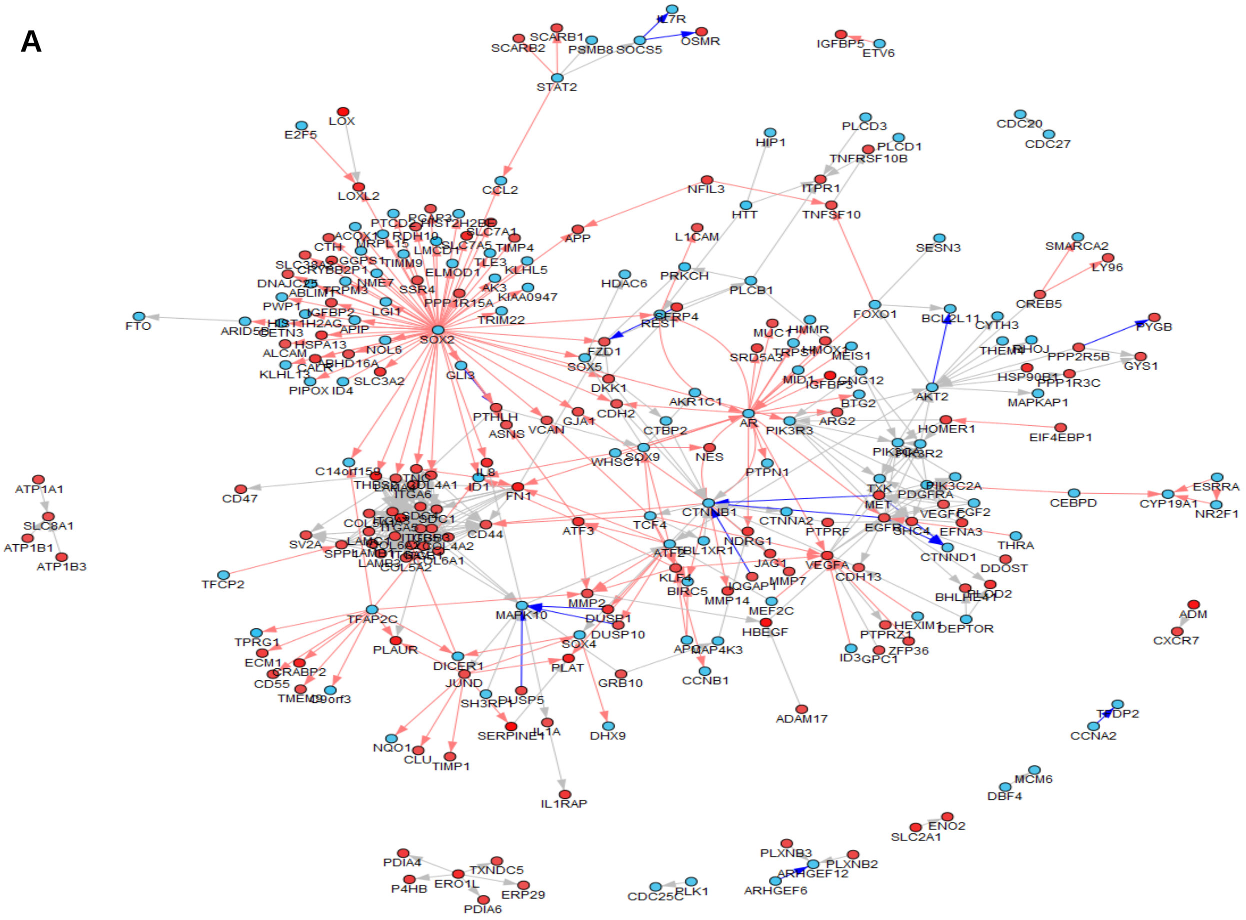


**Supplementary Figure 3: Main DEG network after Mn exposure.** A) Network visualization of significantly affected genes, revealing IL-1A and IL-8 connection to JNK pathway.

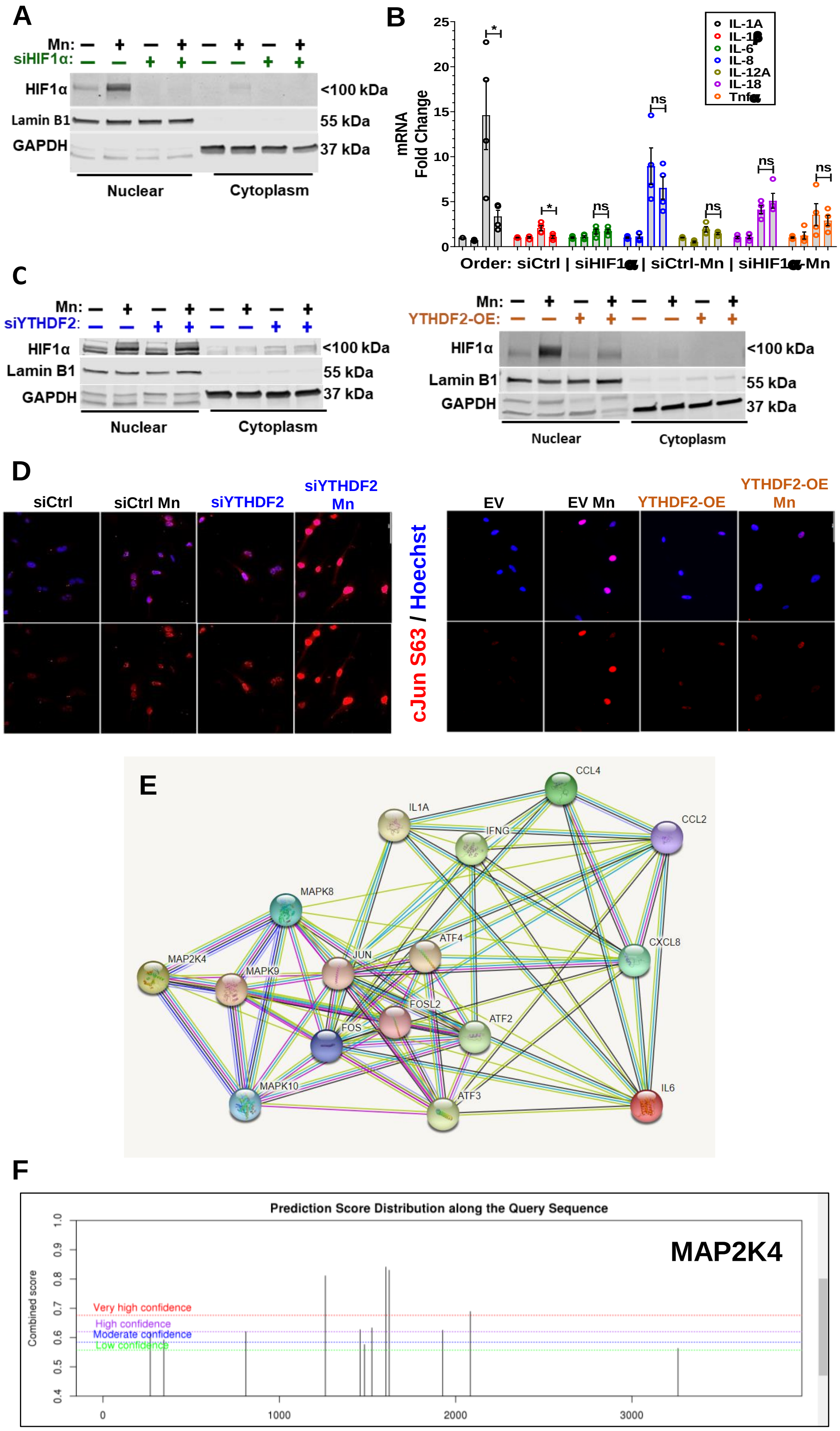
A
Mn:
siHIF1α:
HIF1α
Lamin B1
GAPDH
<100 kDa
55 kDa
37 kDa
Nuclear
Cytoplasm
B
mRNA
Fold Change
IL-1A
IL-6
IL-8
IL-12A
IL-18
ns
Order: siCtrl | siHIF1 | siCtrl-Mn | siHIF1 -Mn
C
siYTHDF2:
YTHDF2-OE:
D
siCtrl
siCtrl Mn
siYTHDF2
siYTHDF2 Mn
cJun S63 / Hoechst
EV
EV Mn
YTHDF2-OE
YTHDF2-OE Mn
E
CCL4
IL1A
IFNG
CCL2
MAPK8
ATF4
CXCL8
JUN
MAP2K4
MAPK9
FOSL2
ATF2
FOS
MAPK10
ATF3
IL6
F
Prediction Score Distribution along the Query Sequence
MAP2K4
Combined score
Very high confidence
High confidence
Moderate confidence
Low confidence

**Supplementary Figure 4: Mn exposure affects both SEK1(*MAP2K4*)-JNK-cJUN pathway and HIF1α in astrocytes.** A) Mn exposures increases HIF1α, but si*HIF1α* abolished the response. B) si*HIF1α* attenuates only *IL-1A* and *IL-1β* gene expression induced by Mn exposure (n=4). C) HIF1α levels after YTHDF2 knockdown and overexpression, in which YTHDF2 overexpression suppressed the level of HIF1α in Mn exposed astrocytes. D) ICC representation at 40x depicting cJUN phosphorylation is increased in Mn exposed astrocytes and sustained in siYTHDF2 Mn exposed astrocytes, but prevented in YTHDF2 overexpressing Mn exposed astrocytes. E) STRING analysis depicting annotated evidence of the SEK1(*MAP2K4*)-JNK-cJUN pathway connected to chemokines/cytokines such as IL-8. F) MAP2K4 m6A sites identified by sRAMP. Data are means ± SEM. Two-way ANOVA with FDR Two-stage step-up method of Benjamini, Krieger and Yekutieli for multi-group comparison. Q-values ≤0.05 considered significant evidence.
